# Site wind right: Identifying low-impact wind development areas in the Central United States

**DOI:** 10.1101/2020.02.11.943613

**Authors:** Chris Hise, Brian Obermeyer, Marissa Ahlering, Jessica Wilkinson, Joseph Fargione

## Abstract

To help avoid the most catastrophic effects of climate change, society needs to achieve net-zero greenhouse gas emissions by mid-century. Wind energy provides a clean, renewable source of electricity; however, improperly sited wind facilities pose known threats to wildlife populations and contribute to degradation of natural habitats. To support a rapid transition to low-carbon energy while protecting imperiled species, we identified potential low-impact areas for wind development in a 17-state region of the central U.S. By combining maps of sensitive habitats and species with wind speed and land use information, we demonstrate that there is significant potential to develop wind energy in the Great Plains while avoiding significant negative impacts to wildlife. These low-impact areas have the potential to yield approximately 1,099-1,832 GW of wind capacity. This is equivalent to 10-18 times current U.S. installed wind capacity. Our analysis demonstrates that ambitious low-carbon energy goals are achievable on sites with minimal risk of wildlife conflict.

## Introduction

A dramatic shift to renewable energy is necessary to limit global warming to 1.5°C and avoid the major threats from climate change [1,2]. Notably, 389 species of birds in the United States are estimated to have an increased risk of vulnerability due to climate change [3]. This change in energy sources will require rapid transitions in global energy production and land use by mid-century [2]. Meeting ambitious targets for greenhouse gas (GHG) emission reductions in the U.S. necessitates a large increase in renewable energy [4–6]. Fortunately, state policy and markets are supporting the buildout of renewable energy. Thirty-eight states have either renewable energy standard portfolios or voluntary targets [7]. The price of renewable energy has fallen dramatically in the past decade and building wind and solar are now often the cheapest way to increase energy production [8].

While wind energy is abundant, renewable, and economic, siting can be challenging. Utility-scale wind facilities require much larger areas of land than conventional forms of electricity generation [9], and renewable energy projects can have significant negative impacts on wildlife and high-priority conservation habitats [10,11]. One concern is direct mortality for migrating bats and birds. Turbines kill hundreds of thousands of bats each year [12]. Direct mortality of songbirds is lower, but still accounts for well over 100,000 bird deaths annually across the U.S. and Canada [13,14]. This is a relatively small number of birds killed compared to other sources of direct mortality from cats, buildings, automobile collisions, power line collisions, power line electrocutions, and communication tower collisions [15]. However, direct mortality from collision with wind turbines may be high enough to have population-level impacts on several species, including migratory tree-roosting bats and golden eagles [11,12,16]. For example, a recent study suggests that the hoary bat (*Lasiurus cinereus*) population of North America could decline by as much as 90% in the next 50 years at current wind energy-associated fatality rates [17].

Impacts of wind turbines on habitat fragmentation and species avoidance is arguably less well appreciated, but of greater conservation concern. Wind turbines also displace wildlife [18] and impact breeding densities of already declining grassland birds [19–21] and waterfowl [22]. Furthermore, the Great Plains, which is home to most of the nation’s remaining intact grasslands, is likely to be one of the areas with the greatest wind energy expansion in the U.S. [9]. Fortunately, many of the ecological concerns with wind energy may be reduced or eliminated by proper siting [11,23,24]. Thus, conservation of birds and bats will require that we transition from fossil fuels to renewable energy production and that we do so in ways that avoid impacts to the most important habitat and employ the best available science to then minimize impacts to habitat and from direct mortality.

Outside of the ecological siting considerations, the stochastic nature of the wind resource itself, along with other anthropogenic and engineering challenges, add further constraints to appropriate siting for wind energy. Wind speed is highly variable on both spatial and temporal scales, which poses challenges for the power grid and for energy delivery [25,26]. Placement of wind development in areas that achieve a certain threshold of wind power throughout the year and have access to transmission is important. Furthermore, areas already developed for urban uses, airports, weather radar, and military training preclude wind development [27–29]. Finally, the height and weight of the turbines limits wind development to mild slopes and stable substrate [29], although technological advances are driving down the cost of wind and now allowing for the development of wind capacity in areas that would have previously not been deemed economically viable [26,30]. Thus, there are a range of ecological, technical, and land use constraints that need to be considered in the siting of wind development.

The mitigation hierarchy provides a framework for development that addresses conservation concerns by first avoiding impacts to the most ecologically important places, and then minimizing impacts through operational practices and offsetting remaining residual impacts through compensatory actions such as habitat protection or restoration [31]. The effectiveness of the mitigation hierarchy is predicated on avoiding impacts to habitat and species to the maximum extent practicable before minimization and offsets are considered. The U.S. Fish and Wildlife Service’s Wind Energy Guidelines (WEGs) [32] lays out a tiered decision framework for evaluating a site for potential wind development that acknowledges the importance of adhering to the hierarchy. The first two tiers of the guidelines recommend evaluating landscape-level considerations before assessing micro-siting, minimization, and offsets. The goal of landscape-level siting is to avoid “potential adverse effects on species of concerns and their habitats” [32], although the WEGs do not specify the areas that need to be avoided. Previous regional efforts to map low-conflict areas for wind energy have focused on identifying disturbed lands where wind could be located with minimal or no habitat concerns [24] in regions outside of the Great Plains [33], or in a subset of the Great Plains [18,34]. This effort advances and significantly expands the geographic scope of previous work.

We identified areas in the central U.S. across 17 states where wind projects are less likely to encounter significant wildlife-related conflict and associated project delays and cost overruns. We used the best available information on sensitive species, natural habitats, and migration corridors that may be adversely impacted by wind development. To provide a realistic estimate of where low impact wind power may be built and to demonstrate its potential to meet the need for low-carbon energy, we also removed areas with low potential for wind development due to engineering constraints and land use conflicts. The resulting maps can be used by power purchasers, developers, and other stakeholders to support landscape-level site screening early in the project planning process.

## Methods

The study area included a 17-state region of the central U.S. (generally conterminous with the Great Plains bioregion) that encompasses 80 percent of the country’s current and planned onshore wind capacity [35]. We chose this geography because wind is rapidly being developed across the landscape, with potentially significant impacts on natural habitats [9].

Building upon previous research [18,24,34], we compiled spatial data on intact natural habitats, imperiled species, and other areas of significant conservation value that may be vulnerable to wind energy impacts. We made use of over 100 sources of data on sensitive habitat, species occurrence, protected areas, and land use. Recognizing that many other siting considerations exist, we also mapped areas with engineering and land use constraints which may render wind development impractical, as identified in published assessments of renewable energy potential and consistent with historical patterns of wind development in the Great Plains. Component spatial data layers are described below.

### Whooping Crane Stopover Sites

The federally endangered whooping crane (*Grus americana*), which has a current population of approximately 500 individuals, depends on wetlands in the central Great Plains during migration [36]. Whooping cranes may be at risk of turbine collisions when ascending or descending from high altitude migration flights, or when travelling short distances between roost and foraging areas [37]. To address this concern, we delineated wind development avoidance areas within 3.2 km of whooping crane stopover sites (Fig 1A). Stopover sites include locations with two or more confirmed whooping crane observations since 1985 [38], as well as modeled suitable habitat [cf. 40–42] within portions of the migratory flyway frequently used by whooping cranes. Modeled suitable habitat included contiguous areas >10 ha in size that met all the following criteria: <100 m from a non-forested, non-rocky wetland or perennial stream [41] or playa lake [42], <1 km from cropland [43], <3% primary and secondary road land cover [44] within a 1 km^2^ moving window, <10% urban land cover [45] within a 1 km^2^ moving window, and intersected core intensity or extended use core intensity areas within a defined migration corridor [46]. We also mapped critical habitat designated by the U.S. Fish and Wildlife Service [47], and whooping crane priority landscapes in Nebraska [48].

**Figure 1.**
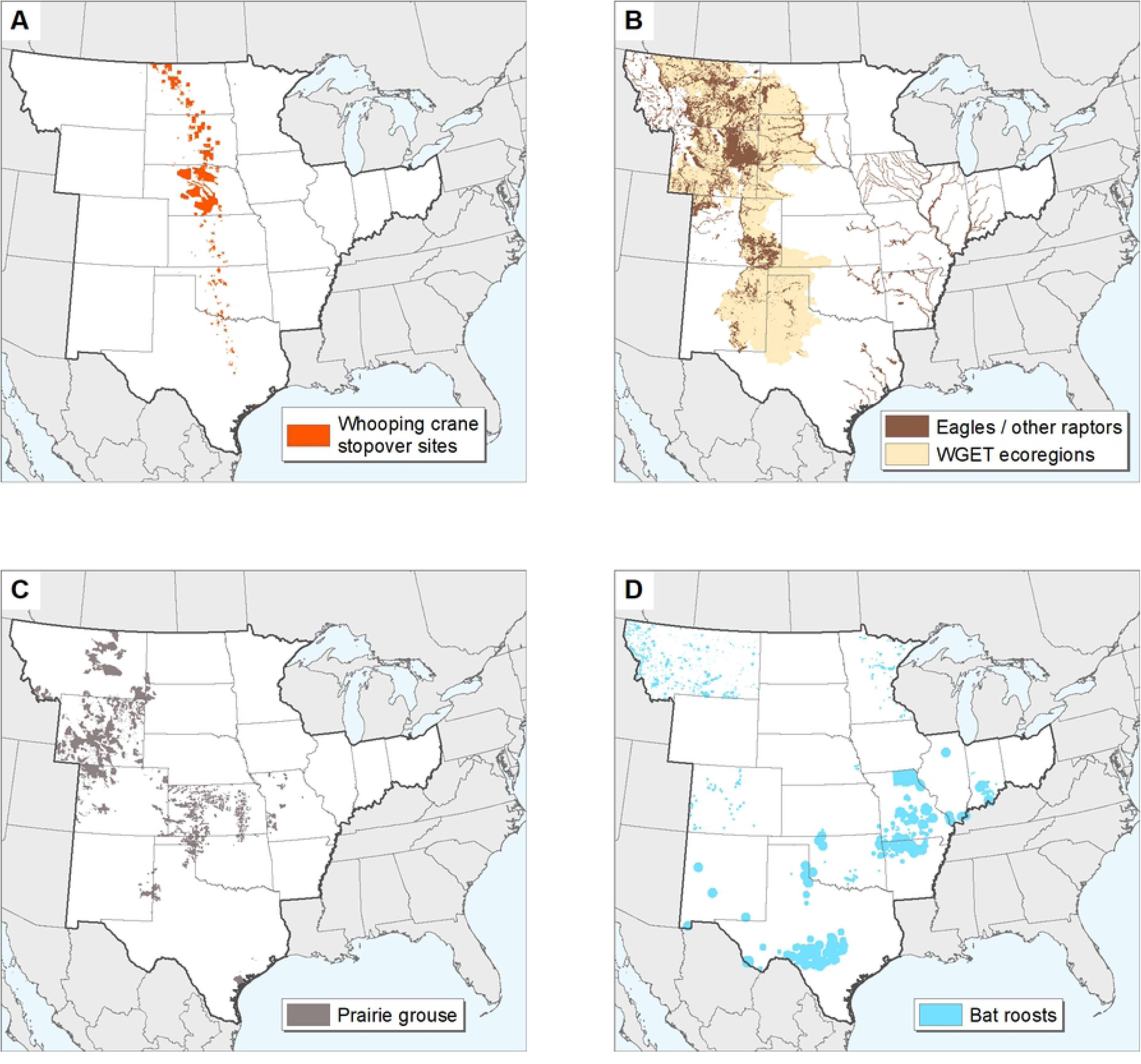
Key areas for birds and bats in the central U.S. (A) Whooping crane stopover sites. (B) High density nesting areas for eagles and other raptors. Brown depicts recommended avoidance areas and tan indicates ecoregions assessed by the U.S. Fish and Wildlife Service Western Golden Eagle Team. (C) Prairie grouse priority habitats. (D) Important bat roosts.

### Eagles and Other Raptors

Raptors may be injured or killed by collisions with wind turbines [49–51], and rates of mortality at commercial wind facilities may be underestimated due to lack of rigorous monitoring and reporting [16]. To reduce the risk of population-level impacts to golden eagles (*Aquila chrysaetos*) in the western Great Plains, we mapped wind development avoidance areas corresponding to the highest modeled golden eagle densities in ecoregions assessed by the Western Golden Eagle Team [53] (top 2 of 7 area-adjusted frequency quantiles; Fig 1B). Following general habitat management guidelines established by the U.S. Fish and Wildlife Service [53], we also mapped avoidance areas within 1.6 km of streams and lakes with known high densities of bald eagle (*Haliaeetus leucocephalus*) nests [54]. In states with available data, we delineated 3.2 km buffers of active golden eagle nests [55] and occupied peregrine falcon (*Falco peregrinus*) habitat [56,57], and 1.6 km buffers of other active raptor nests [55], raptor occurrences [58], and modeled prairie dog (*Cynomys spp.*) complexes due to their attraction of birds of prey [59–62].

### Prairie Grouse

Grouse species in the central U.S. have experienced substantial population declines since the early 20th century [63] and may be further threatened by improperly sited energy development [64–68]. To prevent grouse displacement and potential impacts on vital rates, we mapped the following as important areas to avoid wind development (Fig 1C): Attwater’s prairie-chicken (*Tympanuchus cupido attwateri*) known occurrence records [69] and the Refugio-Goliad Prairie Conservation Area in Texas [70]; Columbian sharp-tailed grouse (*T. phasianellus columbianus*) production areas and winter range in Colorado [71], and 5 km buffers of known leks in Wyoming [72]; greater prairie-chicken (*T. cupido*) modeled optimal habitat [18,54] in Kansas and Oklahoma, production areas in Colorado [73], and grassland conservation opportunity areas in Missouri [74]; greater sage-grouse (*Centrocercus urophasianus*) rangewide biologically significant units [75], state-designated core and connectivity areas in Wyoming [76] and Montana [77], and 2 km buffers of known leks in Wyoming [78]; Gunnison sage-grouse (*C. minimus*) critical habitat [47], and production areas, brood areas, winter range, and severe winter range in Colorado [79]; lesser prairie-chicken (*T. pallidicinctus*) rangewide conservation focal areas and 5 km buffers of known leks [80]; plains sharp-tailed grouse (*T. phasianellus jamesi*) production areas in Colorado [81], and 5 km buffers of known leks in Wyoming [82].

### Bat Roosts

Bat mortality has been documented at wind energy facilities across North America [83–86]. Because bats concentrate in large numbers and have low reproductive rates, the viability of their populations is particularly vulnerable to adult mortality events [87]. Therefore, caution is warranted when undertaking any activity that may adversely affect known bat populations.

While knowledge of bat and wind turbine interactions in the southern Great Plains is limited, evidence suggests that the Mexican free-tailed bat (*Tadarida brasiliensis*) may be particularly susceptible to fatal injury during encounters with turbine blades. This species accounts for a large percentage of documented wildlife mortality at wind facilities across the southwestern U.S. [86,88–90], including in states with extensive wind development. Moreover, regional populations are comprised primarily of reproducing females [91,92]; as such, each early season fatality in the area may result in the death of two individuals (mother and young). Recent population estimates in Oklahoma are markedly lower than historical figures, although the relative contribution of wind is unknown [93]. Due to the large foraging range of this species [94] and concerns regarding population-level impacts, we mapped avoidance areas that extended 32 km from Mexican free-tailed bat maternity roosts [54,95] in New Mexico, Oklahoma, and Texas, as well as adjacent areas of Kansas (Fig 1D).

In addition, we followed the U.S. Fish and Wildlife Service’s [96] recommendation to avoid wind development within 32 km of Indiana bat (*Myotis sodalis*) priority 1 hibernacula, 16 km for priority 2 hibernacula, and 8 km for other current and historical sites. We applied the same rationale and avoidance distances to gray bat (*Myotis grisescens*) hibernacula and other known cave bat roosts across the analysis area [54,97–102]. We also included avoidance areas for mapped bat roosts in Montana [103], mapped hibernacula in Nebraska [48], townships with documented northern long-eared bat (*Myotis septentrionalis*) maternity roosts and/or hibernacula in Minnesota [104], a 12-county region of northeastern Missouri near Sodalis Nature Preserve [105], and important forest habitats in Indiana [106].

### High Waterfowl Breeding Density

We included areas with high predicted waterfowl breeding pair density (15.4 pairs per km^2^ or greater in North Dakota and South Dakota, 9.1 pairs per km^2^ or greater in Minnesota) based on U.S. Fish and Wildlife Service Habitat and Population Evaluation Team data as key areas to avoid wind development (Fig 2A). This represents crucial habitat for many species of wetland-dependent birds in the Prairie Pothole region of North America [107,108].

**Figure 2.**
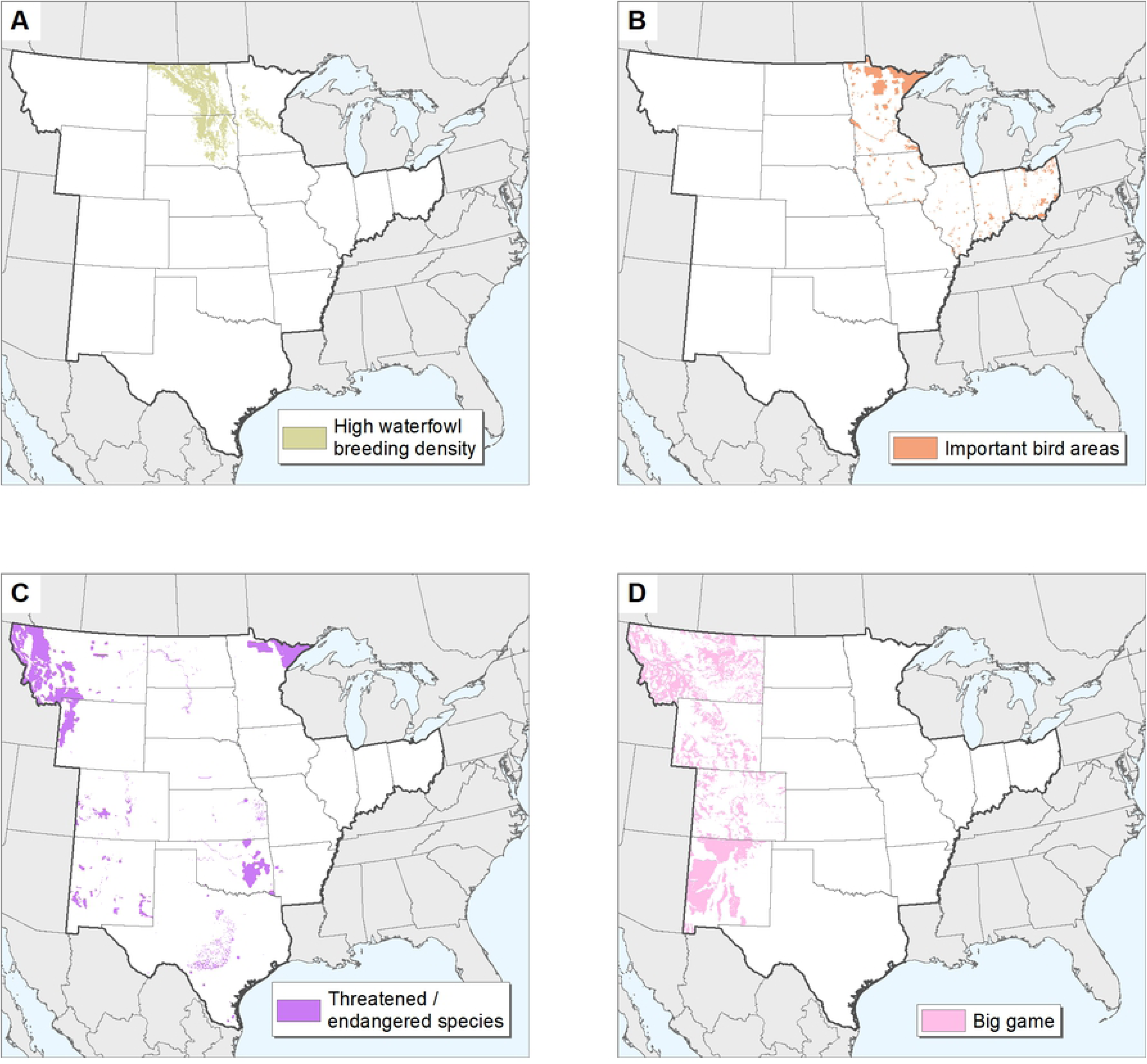
Key areas for birds, threatened and endangered species, and big game in the central U.S. (A) Areas with high predicted waterfowl breeding pair density in the northern Great Plains. (B) Important bird areas in Illinois, Indiana, Iowa, Ohio, and Minnesota. (C) Terrestrial threatened and endangered species habitats. (D) Big game migration corridors and crucial winter range in the Rocky Mountain states.

### Important Bird Areas

Important bird habitats across the Great Lakes and Upper Midwest states may not be effectively captured by other spatial data layers used in this assessment. Therefore, we included state bird conservation areas in Iowa [109] and Audubon important bird areas in Illinois, Indiana, Minnesota, and Ohio [110] as areas to avoid wind development (Fig 2B).

### Other Terrestrial Threatened and Endangered Species

Energy and infrastructure development are among the most significant threats to imperiled species in the U.S. [111]. To prevent impacts to at-risk wildlife, we included terrestrial federally listed threatened and endangered species habitat as avoidance areas. Mapped sites included essential and critical habitat delineated by state and federal agencies [47,112,113], current/recent species distributions [114–118], modeled priority habitats [79,119–121], and occurrence records [54,122,123] (Fig 2C).

### Big game

Roads and other anthropogenic features associated with energy development may alter the movement of big game animals and increase rates of mortality, particularly along migration routes and in crucial winter range in the western U.S. [124–128]. Based on the potential for loss and fragmentation of these vital habitats, we delineated wind development avoidance areas for big game in Montana, Wyoming, Colorado, and New Mexico using available spatial data [129–132] (Fig 2D).

### Important Wetlands and Rivers

Wind energy development near wetland complexes and riparian corridors may cause adverse impacts to migratory birds and other wildlife [18,133–135]. Significant wetland features identified by TNC and partners to avoid wind development included playa clusters and large isolated playa lakes [42]; 1.6 km buffers of important rivers in North Dakota and South Dakota [34], Nebraska [48], Minnesota [136], and Ohio [133]; 16 km buffers of Western Hemisphere Shorebird Reserve Network [137] wetland sites in Illinois, Kansas, Missouri, Oklahoma, and Texas plus the Aransas and Washita National Wildlife Refuges [139, following 35]; and wetlands of special significance [139] and trumpeter swan (*Cygnus buccinator*) occurrence records [140] in Montana (Fig 3A).

**Figure 3.**
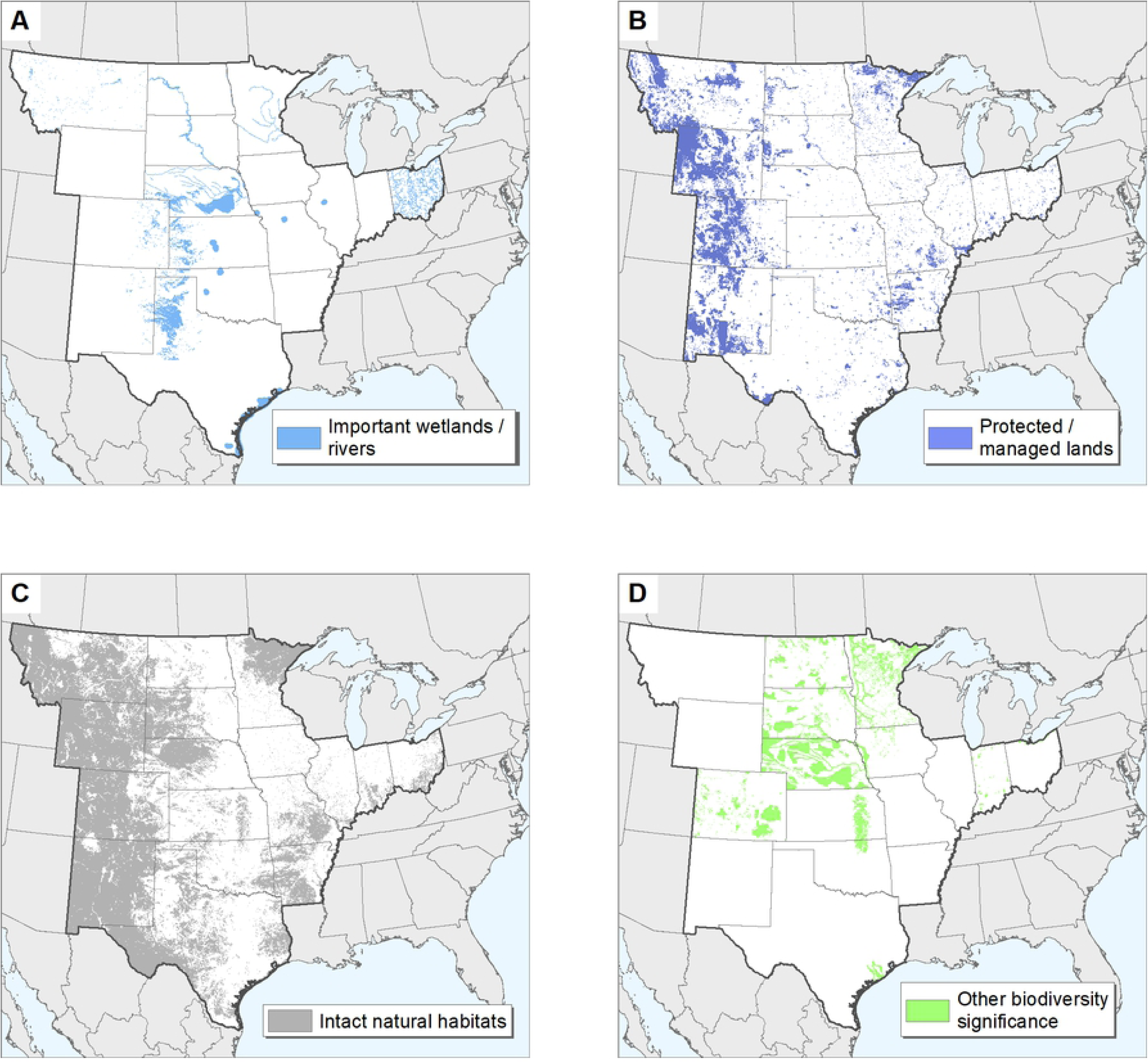
Key lands and waters for wildlife in the central U.S. (A) Important wetlands and rivers. (B) Protected and managed lands. (C) Intact natural habitats. (D) Other areas of biodiversity significance.

### Protected and Managed Lands

We mapped wind development avoidance areas in locations managed for long-term conservation of natural features, including state parks and wildlife management areas; national monuments, parks, and wildlife refuges; military installations; other state and federal lands with development restrictions; private protected lands; and conservation easements [54,138,141]. Due to the relative scarcity and high conservation value of federal lands in the eastern portion the study area, all U.S. Forest Service properties outside of Colorado, Montana, New Mexico, and Wyoming were included regardless of planning designation status (Fig 3B).

### Intact Natural Habitats

Agricultural conversion and other land use change across the central U.S. has significantly reduced the spatial extent of prairie ecosystems and contributed to the loss of many associated species [142]. Remaining intact habitats provide the basis for long-term viability of many species of conservation concern. To delineate discrete patches of undisturbed natural landcover for wind development avoidance, we processed the Theobald [143] human modification (HM) model using a 1 km radius moving window and selected areas with HM index values less than 0.125. We then eliminated areas fragmented by oil and natural gas development, defined as sites with 1.5 active wells per km^2^ or greater [55,145–156, see 127]. We also excluded lands in the Great Plains bioregion altered by past tillage or other landscape disturbances [156]. Finally, we added core forest and core wetland areas [157] to capture additional, functionally intact habitats in Illinois, Indiana, Iowa, Minnesota, Missouri, and Ohio (Fig 3C).

### Other Areas of Biodiversity Significance

We mapped wind development avoidance areas in other areas of recognized conservation importance, including the Flint Hills landscape of Oklahoma and Kansas [137,158]; potential conservation areas with high, very high, or outstanding biodiversity significance in Colorado [159]; wind sensitive areas in Indiana [106]; Prairie Pothole Joint Venture priority areas, and the Loess Hills ecoregion in Iowa [160]; areas of moderate, high, or outstanding biodiversity significance [161], and prairie conservation core areas, corridors, matrix habitat, and strategic habitat complexes in Minnesota [162]; biologically unique landscapes, and medium and high sensitivity natural communities in Nebraska [48]; TNC’s conservation priority areas in North Dakota and South Dakota [34]; areas within 8 km of the Lake Erie shoreline in Ohio [133]; and the Columbia Bottomlands Conservation Area in Texas [163,164] (Fig 3D).

### Airfields

Commercial wind turbines require undisturbed airspace for operation and may present hazards to air travel. Areas within 3 km of public use and military airfield runways [165] were considered unsuitable for wind development [29].

### Special Use Airspace

Special use airspace areas managed by the Federal Aviation Administration contain unusual aerial activity, generally of a military nature. These include ‘alert’ areas which experience high volumes of training flights, ‘restricted’ areas near artillery firing ranges, and ‘prohibited’ areas with significant national security concerns [166]. Placement of wind turbines within these areas may create hazardous flight conditions and compromise military readiness [167]. We considered alert, restricted, and prohibited airspace [168,169] unsuitable for wind energy development. Because these areas are not fully mapped and may change over time, consultation with the U.S. Department of Defense is still advised prior to constructing wind turbines within defined military operating areas, near low-level flight paths, and in areas that may penetrate defense radar lines of sight.

### Radar Stations

Wind turbines may cause interference with radar signals when sited near weather stations and military installations [167,170]. The National Oceanic and Atmospheric Administration (NOAA) requests that developers avoid constructing wind turbines within 3 km of NEXRAD radar installations [171,172]. A larger avoidance distance of 9.26 km was assumed for Department of Defense radar sites [28]. Outside of these areas, mitigation may be required for wind turbines that penetrate radar lines of sight, particularly for structures within 36 km [172].

### Developed Areas

Urban lands [45] and other developed areas [43] were considered unsuitable for commercial wind development [29].

### Existing Wind Facilities

Areas within 1.6 km of existing wind turbines [173] were considered unsuitable for new wind development. This distance represents the typical spacing of turbine strings oriented perpendicularly to prevailing winds in the Great Plains.

### Excessive Slope

Steeply sloping terrain [174] may significantly increase capital costs associated with turbine construction. Areas of slope exceeding 20% were considered unsuitable for wind development [29].

### Water and Wetlands

Open water and wetland areas [41–43] were considered unsuitable for wind development [29].

### Poor Wind Resource

For purposes of this assessment, we considered areas with annual average wind speeds of less than 6.5 m/s at 80 m height to be unsuitable for wind development [175].

### Negative Relative Elevation

Mesoscale wind maps are often generalized and may not accurately depict wind energy potential at a given site [176,177]. Wind developers employ a variety of computational models to assess local wind resources based on orography, measured wind speed, and other factors [178,179]. Most commercial wind facilities in the central U.S. are situated on topographic ridges that experience higher winds than the general surroundings. To identify terrain conducive to development, we calculated relative elevation based on the mean elevation of annuli extending 3, 6, 12, and 24 km from a given point [180]. Negative values represent areas that lie below the adjacent landscape and thus have decreased wind exposure and low potential for wind energy development. Mountainous and coastal regions were not analyzed or excluded based on relative elevation as wind resources in these areas may be influenced by more complex topographic and meteorological factors.

### Statutory Restrictions

Wind development may be legally (or functionally) restricted in some areas of the central U.S., including those within 2.8 km of airport runways, public schools, and hospitals in Oklahoma [165,181–183]); the “Heart of the Flint Hills” region in Kansas [184,185]; 1.6 km and 800 m buffers of certain state-protected properties in Illinois, as supported by the Illinois Natural Areas Preservation Act [186,187]; and 150 m buffers of state trails in Minnesota [188,189]. Many additional state, county, and local regulations pertaining to wind development may exist across the region; however, a detailed examination of these constraints was beyond the scope of this assessment.

### Data Processing

Input data were rasterized at a ground sample distance of 30 m. We generated a preliminary Boolean map of areas suitable for wind development by excluding lands with potential engineering and land use restrictions. To eliminate isolated areas too small to support commercial wind development projects, the results were smoothed using a 1 km radius moving window, and patches less than 20 km^2^ in size were removed. The component engineering and land use restriction layers were then subtracted from the remaining smoothed patches to eliminate false positive values and other spatial artifacts introduced by the moving window analysis. To delineate suitable wind development areas with low potential for wildlife conflict, wildlife and habitat data layers were subtracted from the preliminary Boolean suitability map, and the analysis was repeated as above. For each state and for the entire study area, we quantified wind development potential on all lands suitable for wind development based on wind power, infrastructure, and engineering constraints, as well as the subset of suitable lands identified as low-impact for wildlife, assuming a conservative nameplate capacity density of 3 - 5 MW/km^2^ [29,190]. Based on spacing requirements and current turbine designs, nameplate capacity averages about 5 MW/km^2^. However, installed wind farm developments use an average of 3 MW/km^2^ for projects. This lower bound includes projects that may not still expand to add turbines; once fully built out, these projects may achieve greater land use efficiencies. We note that continuing advances in wind technology may enable even greater efficiencies than the range considered here [e.g., 29].

## Results

After considering factors that may constrain wind development, such as low wind speed and steep terrain, and excluding previously developed areas, statutory setbacks, and unsuitable land use (Fig 4), as well as small and isolated sites, approximately 90 million ha of land or 23% of the total 17 state region was deemed suitable for wind development (Table 1). Given its size, Texas had the highest total ha of suitable area, and Nebraska had the greatest percent of the state considered suitable for wind development. Arkansas has the lowest total area and percent of the state suitable for wind development. If all the suitable land across the study area was developed for wind energy, it could support approximately 2,693 – 4,488 GW of wind capacity.

**Figure 4.**
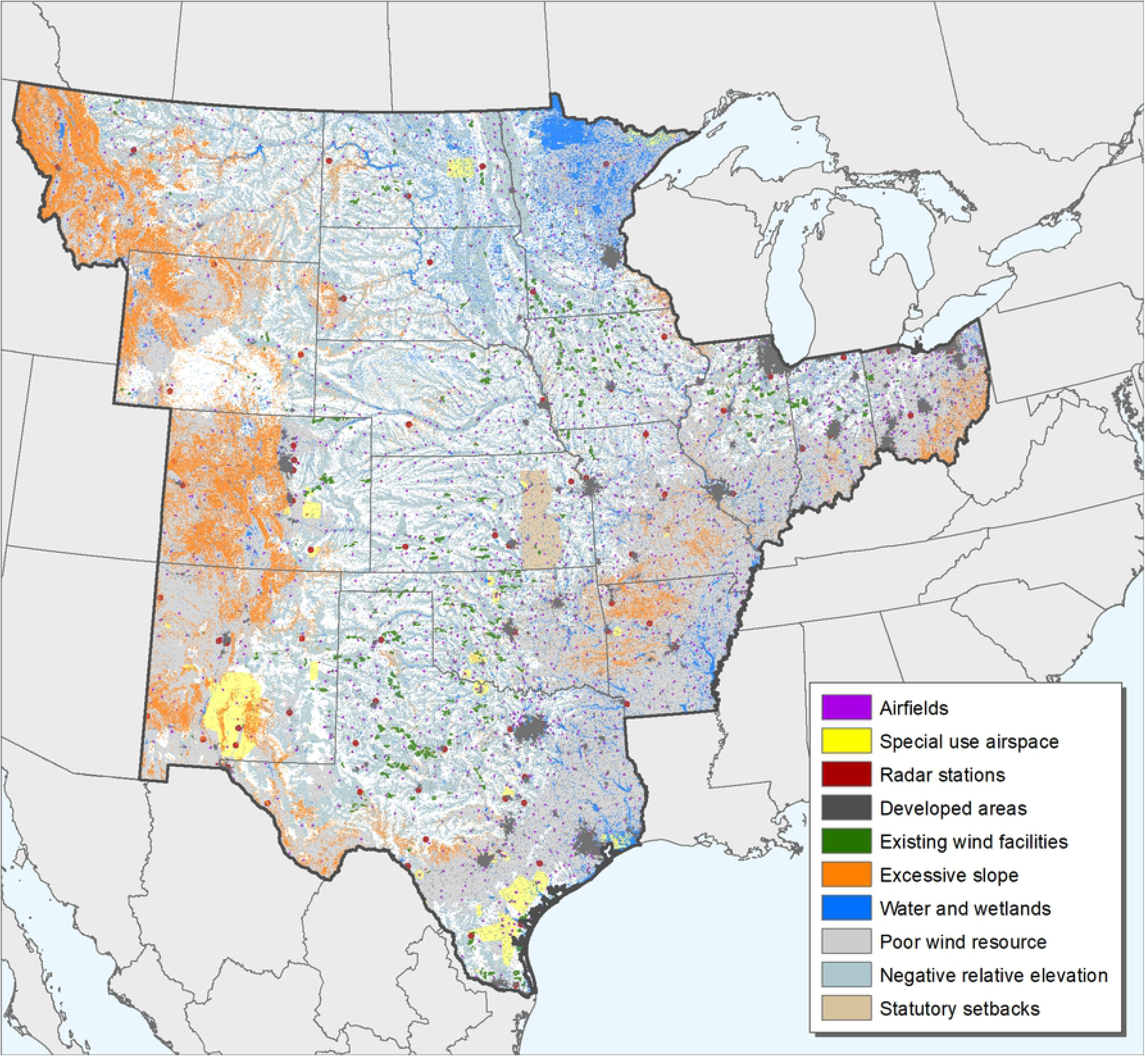
Potential engineering and land use restrictions within a 17-state area of the central U.S.

**Table 1.**
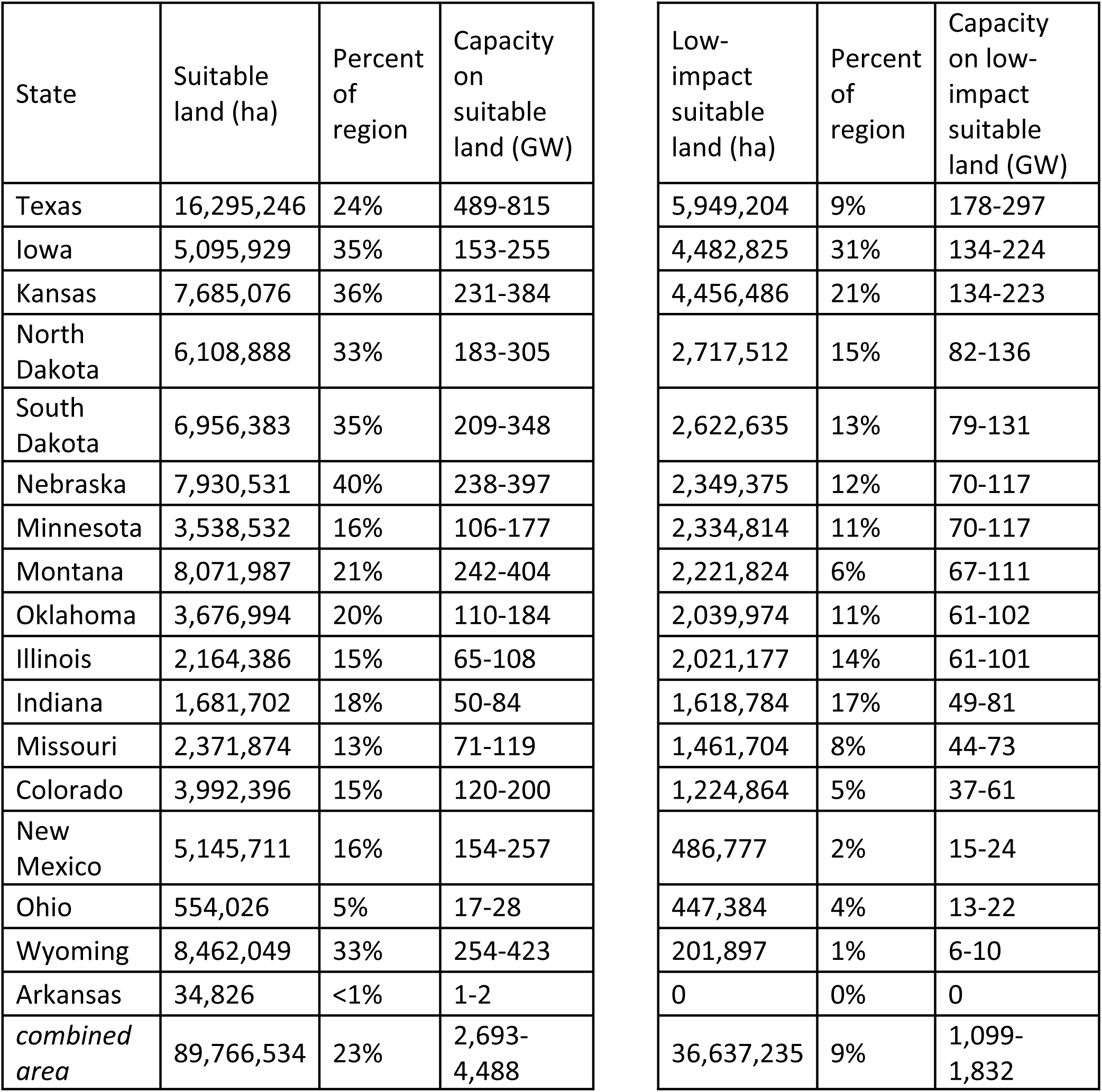
Amount of suitable and low-impact land for wind development and potential wind energy capacity within a 17-state area of the central United States.

After removing sensitive wildlife habitats (Fig 5), approximately 37 million ha remained as suitable for development, which is 9% of the 17-state study area (Fig 6; Table 1). Even though the sensitive wildlife areas mapped across the region included diverse taxa and habitats from big game to bat roosts, there was substantial overlap in where these sensitive wildlife areas occurred. For example, areas mapped as intact natural habitats often co-occurred with sensitive sites such as high-density eagle nesting areas and critical habitat for threatened and endangered species. Once again, Texas had the greatest total area delineated as low-impact to wildlife yet suitable for wind development, comprising 9% of the state. Iowa followed Texas in total amount of low-impact acreage (31% of the state). Nebraska’s low-impact suitable areas included 12% of the state and Arkansas had no low-impact areas deemed suitable for wind development. While Wyoming and New Mexico both have significant wind resources, very little of those states are low-impact for wildlife given the intactness of the habitat. When sensitive wildlife and technical limitations are all considered, we find 1,099 – 1,832 GW of wind capacity potential in low-impact areas alone. This is equivalent to 10 – 18 times current U.S. wind capacity, which reached 100 GW in 2019 [191].

**Figure 5.**
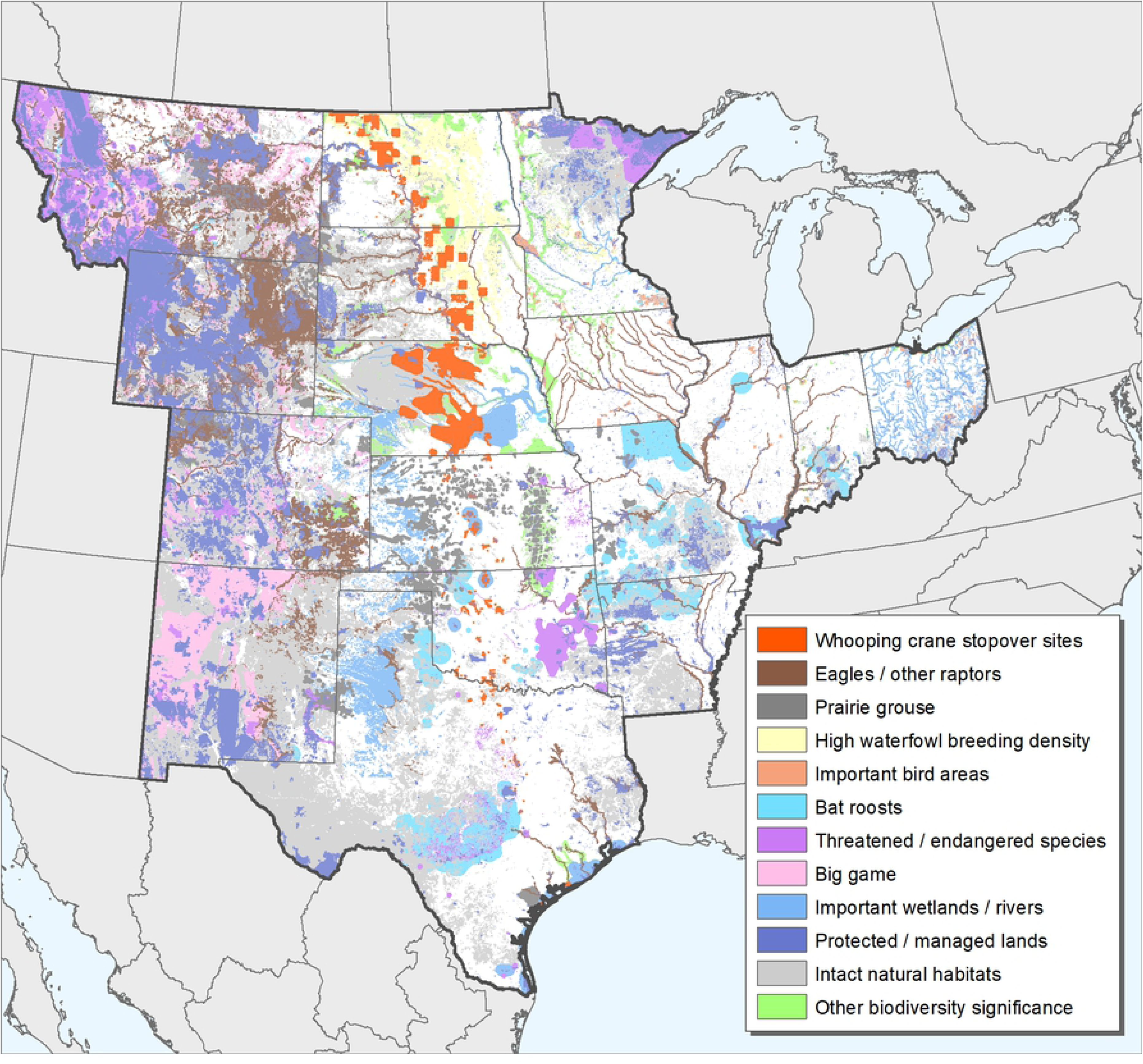
Combined key wildlife areas within a 17-state area of the central U.S.

**Figure 6.**
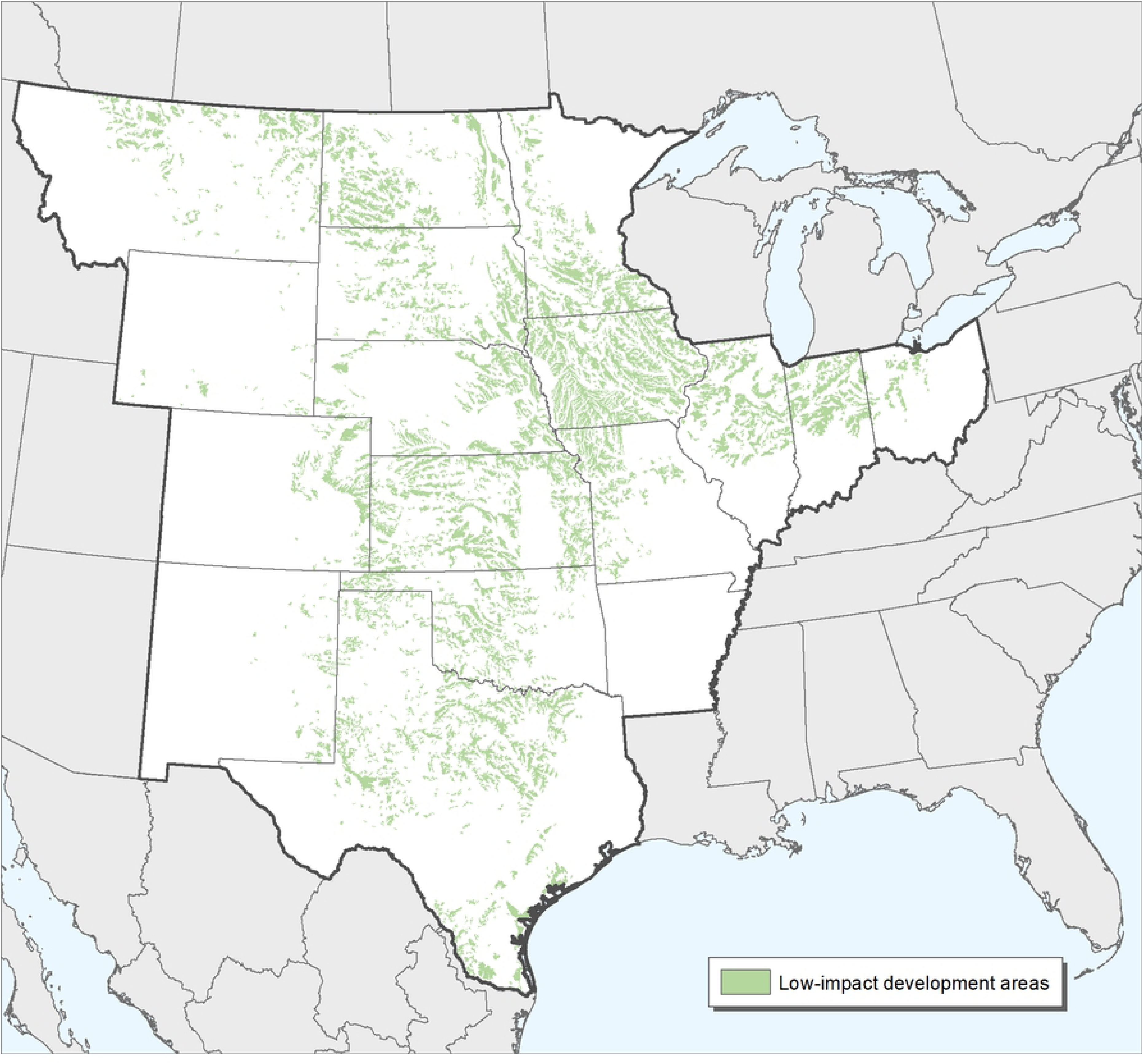
Low-impact areas for wind development within a 17-state area of the central U.S.

## Discussion

Our analysis concludes that there is significant potential to develop wind energy in the central U.S. while avoiding potential impacts to wildlife and sensitive areas. Even in deep decarbonization scenarios where fossil fuels are largely displaced from electricity production, onshore wind energy is expected to generate 386 – 1,392 GW by 2050 [4–6]. This wide range in projected wind energy is due to varying in assumptions about solar and offshore wind, and about overall increases in electricity demand based on electrification of the transportation sector. This means that restricting siting to low-impact areas need not constrain wind development; indeed, even in an unrealistic scenario where all new onshore wind was produced in the study area and grew to produce the large majority of all energy in the United States, there would be adequate low-impact areas to support wind development.

Our analysis demonstrates that ambitious wind development goals are achievable and scalable while minimizing risk to sensitive species and habitats. Our key wildlife areas map (Fig 5) can be used to inform landscape-level siting decisions and can support application of the U.S. Fish and Wildlife Service Wind Energy Guidelines (WEGs) [32], specifically Tier 1 and Tier 2 evaluations. It is not intended to serve as a substitute for the WEGs or to suggest that field surveys or monitoring are not necessary. The map does not replace the need to consider the data and information outlined in the WEGs, consult with state and federal wildlife agencies, or conduct detailed site-level analyses of impacts. Rather, it can be used in conjunction with other appropriate information on habitat and species to support early site screening. To facilitate use of this information by interested parties, we created a web-based mapping application available at http://www.nature.org/sitewindright.

Our estimates of the area suitable for low-impact wind development may be conservative. Some of the low-impact areas we identified as having engineering and land use constraints may be viable for wind energy due to improvements in technology [192]. For example, taller turbines and longer blades are making wind energy development economically viable even in areas with lower wind speeds [30]. Countervailing factors include the potential for transmission capacity and the availability of willing landowners to limit the development of some of the low-impact areas that we identified [28,193]. Additionally, because there are many unknown factors regarding how wind development and the wind turbines themselves impact wildlife [11] we used conservative estimates to identify areas that should be considered for avoidance due to conservation concerns. The impact of wind turbines on direct mortality has received the most attention, but other aspects of wind development such as displacement or the potentially synergistic effects of multiple wind developments on the landscape are not well understood. Some studies have documented displacement effects for grassland birds [19] and waterfowl [22], but how widespread or long-term these impacts are is unclear. Given the relatively large footprint of wind development [9], the decline of many grassland bird species [21], the unknown degree and geographic extent of impacts [11], and the amount of low-impact wind development area available, targeting development in low-impact areas would allow us to both reduce carbon emissions and reduce the risk of impacts to wildlife habitat and populations.

We also note that our delineation of sensitive wildlife habitats was not exhaustive and should be updated as our understanding of the impacts of wind on wildlife continue to improve. Spatial data on species of concern are missing or incomplete in many geographies and for many taxa. For example, seasonal or temporary wetlands have a high value for waterfowl, particularly in the Prairie Pothole Region, but can be difficult to map, especially when embedded in agricultural fields or other disturbed areas [194] and the distribution and migratory pathways of bats are not well known [87,195]. Therefore, as with all development projects, wildlife concerns should be addressed early in the site identification process, should be made in the context of landscape-level considerations, and should be fully explored before considering minimization and offset measures. For some species, operational minimization measures may be necessary to reduce mortality, particularly for bats [196,197]. We strongly encourage the use of proven operational minimization strategies [196] and new approaches such as smart curtailment [198] to reduce impacts to bat populations. Bats appear to be attracted to turbines, contributing to their high mortality rates [199], but curtailing wind production during nights with high migratory activity, increasing turbine cut-in speed [200,201], and the use of ultrasonic deterrents [202] can substantially reduce mortality. New methods to track seasonal bat movements are in development [203] and could further inform operational minimization strategies.

Our map of key wildlife areas may serve as an important source of information for developers to support screening early in the project siting process and for power purchasers and consumers interested in wildlife-friendly wind power. Additionally, power purchasers acquiring wind-generated electricity from low-impact sites may be able to support their climate and renewable energy goals, while also avoiding sensitive species and habitats. Projects proposed in the low-impact areas are less likely to encounter wildlife-related conflict and associated project delays and related cost overruns, which should speed deployment of the wind energy needed to meet climate mitigation goals [28].

## Acknowledgements

We thank many colleagues at The Nature Conservancy who contributed data and insight to this assessment, including (in alphabetical order): Matt Bain, Steven Bassett, Fiona Becker, Viv Bennett, Erica Brand, Ken Brunson, Angela Burke, Dick Cameron, Teresa Chapman, Brian Cohen, James Cole, Holly Copeland, Nathan Cummins, Neal Feeken, Robert Findling, August Froehlich, Michael Fuhr, Steven Gilbert, Jan Glendening, Katie Gillies, Katie Hawk, Sarah Hagen, Bob Hamilton, Fran Harty, Jim Hays, Chris Helzer, Susanne Hickey, Tharran Hobson, Shannon Jauregui, Nels Johnson, Rich Johnson, Clayton Knighten, Rich Kostecke, Aaron Lange, Robert Manes, Brian Martin, Sara Mascola, Ashley Maybanks, Bruce McKenney, Scott Moats, Holly Neill, Marcella Ochoa, Amy Pearson, Cynthia Pessoni, Corinna Riginos, John Sanderson, Melissa Shackford, John Shuey, Jan Slaats, Michael Slay, Bill Stanley, Karla Suckling, Jeffery Walk, Nick Walters, Rich Walters, Rachel Worthen, and Douglas Zollner.

## Author Contributions

Conceived and designed the experiments: CH BO JF. Performed the experiments and analyzed the data: CH. Wrote the paper: CH MA BO JW JF.

## References

1. Tallis HM, Hawthorne PL, Polasky S, Reid J, Beck MW, Brauman K, et al. An attainable global vision for conservation and human well-being. Front Ecol Environ. 2018;16: 563–570. doi:10.1002/fee.1965

2. Rogelj J, Shindell D, Jiang K, Fifita S, Forster P, Ginzburg V, et al. Mitigation pathways compatible with 1.5°C in the context of sustainable development. In: Masson-Delmotte V, Zhai P, H.-O., Pörtner DR, Skea J, Shukla PR, et al., editors. Global warming of 15°C An IPCC Special Report on the impacts of global warming of 15°C above pre-industrial levels and related global greenhouse gas emission pathways, in the context of sustainable development. IPCC; 2018. pp. 93–174.

3. Wilsey C, Bateman B, Taylor L, Wu JX, LeBaron G, Shepherd R, et al. Survival by degrees: 389 species on the brink. New; 2019. Available: https://www.audubon.org/sites/default/files/climatereport-2019-english-lowres.pdf

4. Hand MM, Baldwin S, DeMeo E, Reilly JM, Mai T, Arent D, et al. Renewable electricity futures study. Technical report NREL/TP-6A20-52409. Golden, Colorado; 2012. doi:NREL/TP-6A20-52409-1

5. Williams JH, Haley B, Kahrl F, Moore J, Jones AD, Torn MS, et al. Pathways to deep decarbonization in the United States. The U.S. report of the Deep Decarbonization Pathways Project of the Sustainable Development Solutions Network and the Institute for Sustainable Development and International Relations. Revis with Tech Suppl Novemb 15, 2015. 2015.

6. Haley B, Jones R, Kwok G, Hargreaves J, Farbes J, Williams JH. 350 PPM pathways for the United States. 2019. Available: https://www.evolved.energy/

7. National Conference of State Legislatures. State renewable portfolio standards and goals. 2019 [cited 19 Dec 2019]. Available: http://www.ncsl.org/research/energy/renewable_portfolio_standards.aspx

8. Lazard. Lazard’s levelised cost of energy analysis, version 13.0. 2019. Available: https://www.lazard.com/media/451086/lazards-levelized-cost-of-energy-version-130-vf.pdf

9. McDonald RI, Fargione J, Kiesecker J, Miller WM, Powell J. Energy sprawl or energy efficiency: Climate policy impacts on natural habitat for the United States of America. PLoS One. 2009;4.

10. Arnett EB, Inkley DB, Johnson DH, Larkin RP, Manes S, Manville AM, et al. Impacts of wind energy facilities on wildlife and wildlife habitat. Wildl Soc Tech Rev 07–2. Bethesda, Maryland; 2007.

11. Allison TD, Diffendorfer JE, Baerwald EF, Beston JA, Drake D, Hale AM, et al. Impacts To wildlife of wind energy siting and operation in the United States. Issues Ecol. 2019.

12. Smallwood KS. Comparing bird and bat fatality-rate estimates among North American wind-energy projects. Wildl Soc Bull. 2013;37: 19–33. doi:10.1002/wsb.260

13. Erickson WP, Wolfe MM, Bay KJ, Johnson DH, Gehring JL. A comprehensive analysis of small-passerine fatalities from collision with turbines at wind energy facilities. PLoS One. 2014;9. doi:10.1371/journal.pone.0107491

14. Johnson DH, Loss SR, Shawn Smallwood K, Erickson WP. Avian fatalities at wind energy facilities in North America: a comparison of recent approaches. Human-Wildlife Interact. 2016;10: 7–18.

15. Loss SR, Will T, Marra PP. Direct mortality of birds from anthropogenic causes. Annu Rev Ecol Evol Syst. 2015;46: 99–120. doi:10.1146/annurev-ecolsys-112414-054133

16. Pagel JE, Kritz KJ, Millsap BA, Murphy RK, Kershner EL, Covington S. Bald eagle and golden eagle mortalities at wind energy facilities in the contiguous United States. J Raptor Res. 2013;47: 311–315. doi:10.3356/jrr-12-00019.1

17. Frick WF, Baerwald EF, Pollock JF, Barclay RMR, Szymanski JA, Weller TJ, et al. Fatalities at wind turbines may threaten population viability of a migratory bat. Biol Conserv. 2017;209: 172–177. doi:10.1016/j.biocon.2017.02.023

18. Obermeyer B, Manes R, Kiesecker J, Fargione J, Sochi K. Development by design: mitigating wind development’s impacts on wildlife in Kansas. PLoS One. 2011;6: e26698. doi:10.1371/journal.pone.0026698.

19. Shaffer JA, Buhl DA. Effects of wind-energy facilities on breeding grassland bird distributions. Conserv Biol. 2016;30: 59–71. doi:10.1111/cobi.12569

20. Shaffer JA, Loesch CR, Buhl DA. Estimating offsets for avian displacement effects of anthropogenic impacts. Ecol Appl. 2019;29: 1–15. doi:10.1002/eap.1983

21. Rosenberg K V, Dokter AM, Blancher PJ, Sauer JR, Smith AC, Smith PA, et al. Decline of the North American avifauna. Science (80-). 2019;366: 120 LP–124. doi:10.1126/science.aaw1313

22. Loesch CR, Walker JA, Reynolds RE, Gleason JS. Effect of wind energy development on breeding duck densities in the Prairie Pothole Region. J Wildl Manage. 2013;77: 587–598.

23. Diffendorfer JE, Dorning MA, Keen JR, Kramer LA, Taylor R V. Geographic context affects the landscape change and fragmentation caused by wind energy facilities. PeerJ. 2019;2019: 1–23. doi:10.7717/peerj.7129

24. Kiesecker JM, Evans JS, Fargione J, Doherty K, Foresman KR, Kunz TH, et al. Win-Win for Wind and Wildlife: A Vision to Facilitate Sustainable Development. PLoS One. 2011;6: e17566. doi:10.1371/journal.pone.0017566. doi:e1756610.1371/journal.pone.0017566

25. Bouffard F, Galiana FD. Stochastic security for operations planning with significant wind power generation. IEEE Power Energy Soc 2008 Gen Meet Convers Deliv Electr Energy 21st Century, PES. 2008;23: 306–316. doi:10.1109/PES.2008.4596307

26. Veers P, Dykes K, Lantz E, Barth S, Bottasso CL, Carlson O, et al. Grand challenges in the science of wind energy. Science (80-). 2019;366. doi:10.1126/science.aau2027

27. Lopez A, Roberts B, Heimiller D, Blair N, Porro G. U.S. renewable energy technical potentials: a GIS-based analysis. Technical report NREL/TP-6A20-51946. Golden, Colorado; 2012. doi:NREL/TP-6A20-51946

28. Tegen S, Lantz E, Mai T, Heimiller D, Hand M, Ibanez E. An initial evaluation of siting considerations on current and future wind deployment. Technical report NREL/TP-5000-61750. Golden, Colorado; 2016.

29. U.S. Department of Energy. 20% wind energy by 2030: increasing wind energy’s contribution to U.S. electricity supply. 2008. Available: http://www.nrel.gov/docs/fy08osti/41869.pdf

30. Lantz E, Roberts O, Nunemaker J, Demeo E, Dykes K, Scott G. Increasing wind turbine tower heights: opportunities and challenges. Technical report NREL/TP-5000-73629. Golden, Colorado; 2019.

31. Kiesecker JM, Copeland H, Pocewicz A, McKenney B. Development by design: blending landscape-level planning with the mitigation hierarchy. Front Ecol Environ. 2010;8: 261–266. doi:10.1890/090005

32. U.S. Fish and Wildlife Service. Land-based wind energy guidelines. 2012. Available: https://www.fws.gov/midwest/wind/resources/

33. Evans JS, Kiesecker JM. Shale gas, wind and water: assessing the potential cumulative impacts of energy development on ecosystem services within the Marcellus play. PLoS One. 2014;9. doi:10.1371/journal.pone.0089210

34. Fargione J, Kiesecker J, Slaats MJ, Olimb S. Wind and wildlife in the Northern Great Plains: identifying low-impact areas for wind development. PLoS One. 2012;7.

35. American Wind Energy Association. U.S. wind industry fourth quarter 2018 market report. Washington, D.C.; 2019.

36. U.S. Fish and Wildlife Service. Species profile for whooping crane. 2011 [cited 13 Mar 2015]. Available: http://ecos.fws.gov

37. U.S. Fish and Wildlife Service. Whooping cranes and wind development - an issue paper. 2009. Available: http://www.fws.gov/southwest/es/Documents/R2ES/Whooping Crane and Wind Development FWS issue paper - final April 2009.pdf

38. U.S. Fish and Wildlife Service. Cooperative whooping crane tracking project GIS database. Grand Island, Nebraska; 2010.

39. Austin JE, Richert AL. A comprehensive review of observational and site evaluation data of migrant whooping cranes in the United States, 1943-1999. Jamestown, North Dakota; 2001. doi:10.1146/annurev.polisci.6.121901.085546

40. Belaire JA, Kreakie BJ, Keitt T, Minor E. Predicting and mapping potential whooping crane stopover habitat to guide site selection for wind energy projects. Conserv Biol. 2014;28: 541–550. doi:10.1111/cobi.12199

41. U.S. Fish and Wildlife Service. National wetlands inventory version 2. 2016. Available: http://www.fws.gov/wetlands

42. Playa Lakes Joint Venture. Playa lakes decision support system. 2015 [cited 3 Jul 2018]. Available: http://pljv.org/playa-dss

43. Homer C, Dewitz J, Yang L, Jin S, Danielson P, Xian G, et al. Completion of the 2011 national land cover database for the conterminous United States – representing a decade of land cover change information. Photogramm Eng Remote Sensing. 2015;81: 345–354. doi:10.1016/S0099-1112(15)30100-2

44. ESRI. U.S. and Canada detailed streets. Redlands, California; 2010.

45. U.S. Census Bureau. Urban areas national shapefile (2010 census). 2016. Available: https://www.census.gov

46. Pearse AT, Brandt DA, Harrell WC, Metzger KL, Baasch DM, Hefley TJ. Whooping crane stopover site use intensity within the Great Plains. U.S. Geol Surv Open-File Rep 2015-1166. 2015; 12 pp. doi:doi: 10.3133/ofr20151166

47. U.S. Fish and Wildlife Service. Threatened & endangered species critical habitat polygons shapefile. 2018. Available: https://ecos.fws.gov

48. The Nebraska Wind and Wildlife Working Group. Guidelines for avoiding, minimizing, and mitigating impacts of wind energy on biodiversity in Nebraska. Lincoln; 2016. Available: https://wind-energy-wildlife.unl.edu/

49. Stewart GB, Pullin AS, Coles CF. Poor evidence-base for assessment of windfarm impacts on birds. Environ Conserv. 2007;34: 1–11. doi:10.1017/S0376892907003554

50. Smallwood KS, Thelander C. Bird mortality in the Altamont Pass wind resource area, California. J Wildl Manage. 2008;72: 215–223. doi:10.2193/2007-032

51. Watson RT, Kolar PS, Ferrer M, Nygård T, Johnston N, Hunt WG, et al. Raptor interactions with wind energy: case studies from around the world. J Raptor Res. 2018;52: 1–18. doi:10.3356/jrr-16-100.1

52. Bedrosian G, Carlisle JD, Wallace ZP, Bedrosian B, LaPlante DW, Woodbridge B, et al. Spatially explicit landscape-scale risk assessments for breeding and wintering golden eagles in the western United States. Rep Prep by West Golden Eagle Team, U.S. Fish Wildl Serv Reg 1, 2, 6, 8. 2018.

53. U.S. Fish and Wildlife Service. Southeastern states bald eagle recovery plan. 1989. Available: http://ecos.fws.gov

54. The Nature Conservancy. Oklahoma chapter geographic information system database. Tulsa; 2019.

55. Colorado Parks and Wildlife. Cooperative raptor database. Fort Collins; 2013.

56. Wyoming Game and Fish Department. Occupied peregrine falcon habitat. Cheyenne; 2004.

57. Colorado Parks and Wildlife. Peregrine falcon nesting areas. Fort Collins; 2015.

58. Montana Natural Heritage Program. Species occurrence data for raptors. Helena; 2018.

59. Wyoming Game and Fish Department. White-tailed prairie dog towns. Cheyenne; 2006.

60. Colorado Parks and Wildlife. SPICE habitat model for black-tailed prairie dog. Fort Collins; 2009.

61. Montana Natural Heritage Program. Species occurrence data for white-tailed prairie dog and black-tailed prairie dog. Helena; 2018.

62. Texas Natural Diversity Database. Prairie dog town element occurrence data export. Austin; 2018.

63. Vodehnal WL, Haufler JB. A grassland conservation plan for prairie grouse. Fruita, Colorado; 2007.

64. Pruett CL, Patten MA, Wolfe DH. Avoidance behavior by prairie grouse: implications for development of wind energy. Conserv Biol. 2009;23: 1253–1259. doi:10.1111/j.1523-1739.2009.01254.x

65. Val Pelt WE, Kyle S, Pitman J, Klute D, Beauprez G, Schoeling D, et al. The lesser prairie-chicken range-wide conservation plan. Cheyenne, Wyoming; 2013.

66. Hovick TJ, Elmore RD, Dahlgren DK, Fuhlendorf SD, Engle DM. Evidence of negative effects of anthropogenic structures on wildlife: A review of grouse survival and behaviour. J Appl Ecol. 2014;51: 1680–1689. doi:10.1111/1365-2664.12331

67. Winder VL, Gregory AJ, McNew LB, Sandercock BK. Responses of male greater prairie-chickens to wind energy development. Condor. 2015;117: 284–296. doi:10.1650/condor-14-98.1

68. Lebeau CW, Beck JL, Johnson GD, Nielson RM, Holloran MJ, Gerow KG, et al. Greater sage-grouse male lek counts relative to a wind energy development. Wildl Soc Bull. 2017;41: 17–26. doi:10.1002/wsb.725

69. Texas Natural Diversity Database. Attwater’s prairie-chicken element occurrence data export. Austin; 2018.

70. The Nature Conservancy. Conservation action plan for the Refugio-Goliad Prairie Conservation Area. Austin, Texas; 2009.

71. Colorado Parks and Wildlife. Columbian sharp-tailed grouse production areas and winter range. Fort Collins; 2015.

72. Wyoming Game and Fish Department. Columbian sharp-tailed grouse leks. Cheyenne; 2016.

73. Colorado Parks and Wildlife. Greater prairie chicken production areas. Fort Collins; 2015.

74. Missouri Department of Conservation. Missouri state wildlife action plan. Jefferson City; 2015. Available: https://mdc.mo.gov/

75. Bureau of Land Management. BLM GRSG western U.S. biologically significant units maps service. 2018. Available: https://landscape.blm.gov

76. Wyoming Game and Fish Department. Sage grouse core and connectivity areas version 4. 2015. Available: https://wgfd.wyo.gov

77. Montana Fish Wildlife and Parks. Sage-grouse core areas. 2016. Available: http://gis-mtfwp.opendata.arcgis.com

78. Wyoming Game and Fish Department. Greater sage-grouse leks (occupied). Cheyenne; 2017.

79. Colorado Parks and Wildlife. Gunnison sage-grouse production areas, brood areas, winter range, and severe winter range. Fort Collins; 2011.

80. Kansas Biological Survey. Southern Great Plains crucial habitat assessment tool. 2018 [cited 11 Apr 2018]. Available: http://www.kars.ku.edu/maps/sgpchat

81. Colorado Parks and Wildlife. Plains sharp-tailed grouse production areas. Fort Collins; 2015.

82. Wyoming Game and Fish Department. Plains sharp-tailed grouse leks. Cheyenne; 2017.

83. Erickson W, Johnson G, Young D, Strickland D, Good R, Bourassa M, et al. Synthesis and comparison of baseline avian and bat use, raptor nesting and mortality information from proposed and existing wind developments. Final Rep Submitt to Bonnev Power Adm. Cheyenne, Wyoming; 2002.

84. U.S. Fish and Wildlife Service. Interim guidelines To avoid and minimize wildlife impacts from wind turbines. 2003. Available: http://www.fws.gov/habitatconservation/wind.pdf

85. Arnett E., Baerwald EF. Impacts of wind energy development on bats: implications for conservation. Bat evolution, ecology, and conservation. New York: Springer Science & Business Media; 2013. pp. 435–456.

86. American Wind Wildlife Institute (AWWI). A summary of bat fatality data in a nationwide database. 2018. Available: www.awwi.org

87. Kunz TH, Fenton MB. Bat ecology. University of Chicago Press; 2003.

88. Kerlinger P, Curry R, Culp L, Jain A, Wilkerson C, Fischer B, et al. Post-construction avian monitoring study for the High Winds Wind Power project, Solano County, California: two year report. McLean, Virginia; 2006.

89. Miller A. Patterns of avian and bat mortality at a utility-scaled wind farm on the southern High Plains. Master’s thesis. Texas Tech University, Lubbock. 2008.

90. Piorkowski MD, O’Connell TJ. Spatial pattern of summer bat mortality from dollisions with wind turbines in mixed-grass prairie. Am Midl Nat. 2010;164: 260–269. doi:10.1674/0003-0031-164.2.260

91. Caire W, Tyler JD, Glass BP, Mares MA. Mammals of Oklahoma. Norman: University of Oklahoma Press; 1989.

92. Schmidly DJ. The mammals of Texas (revised edition). Austin: University of Texas Press; 2004.

93. Caire W, Matlack RS, Ganow KB. Population size estimations of Mexican free-tailed bat, Tadarida brasiliensis, at important maternity roosts in Oklahoma. Final Perform report, Fed aid grant F10AF00236. Oklahoma City; 2013.

94. Best TL, Geluso KN. Summer foraging range of Mexican free-tailedc bats (Tadarida brasiliensis mexicana) from Carlsbad Cavern, New Mexico. Southwest Nat. 2003;48: 590–596.

95. Texas Speleological Survey. Known bat caves in Texas (generalized to USGS quadrangle). Austin; 2018.

96. U.S. Fish and Wildlife Service. Midwest wind energy multi-species habitat conservation plan (public review draft). 2016. Available: http://www.midwestwindhcp.com/

97. Kansas State University. Habitat model for gray myotis (Myotis grisescens). 2002. Available: http://www.k-state.edu/kansasgap/KS-GAPPhase1/finalreport/SppModels/Mammals/Gray_Myotis.pdf

98. The Nature Conservancy. Ozarks ecoregional conservation assessment. Minneapolis, Minnesota; 2003.

99. Graening GO, Harvey MJ, Puckette WL, Stark RC, Sasse DB, Hensley SL, et al. Conservation status of the endangered Ozark big-eared bat (Corynorhinus Townsendii Ingens) - a 34-Year assessment. Publ Oklahoma Biol Surv. 2011;11: 1–16.

100. Indiana Natural Heritage Inventory Data Center. Indiana natural heritage database element occurrence records. Indianapolis; 2014.

101. Kansas Biological Survey. Kansas natural resource planner occurrence records for bat caves. 2015. Available: http://kars.ku.edu/maps/naturalresourceplanner/

102. Colorado Natural Heritage Program. Level 3 element occurrence records. Fort Collins; 2017.

103. Montana Natural Heritage Program. Species occurrrence data for bats. Helena; 2018.

104. Minnesota Department of Natural Resources and U.S Fish and Wildlife Service. Townships containing documented northern long-eared bat maternity roost trees and/or hibernacula entrances in Minnesota. 2018. Available: https://files.dnr.state.mn.us

105. Cole J. Notes from a March 7, 2018 meeting with U.S Fish and Wildlife Service staff (Karen Herrington, Shauna Marquardt, and Jane Ledwin) at the ecological services field office in Columbia, Missouri. Jefferson City, Missouri; 2018.

106. The Nature Conservancy and National Audubon Society. Wind power development sensitive areas in Indiana. Indianapolis; 2010.

107. Reynolds RE, Shaffer TL, Loesch CR, Cox RCJ. The Farm Bill and duck production in the Prairie Pothole Region: increasing the benefits. Wildl Soc Bull. 2006;34: 963–974. doi:10.2193/0091-7648(2006)34[963:tfbadp]2.0.co;2

108. Niemuth ND, Reynolds RE, Granfors RR, Johnson RR, Wangler B, Estey M. Landscape-level planning for conservation of wetland birds in the U.S Prairie Pothole region. In: Millspaugh JJ, Thompson FR, editors. Models for planning wildlife conservation in large landscapes. Elsevier Science; 2008. pp. 533–560.

109. Iowa Department of Natural Resources. Bird conservation areas in Iowa. 2017. Available: https://geodata.iowa.gov

110. National Audubon Society. Important bird areas. 2018 [cited 7 Jul 2018]. Available: https://www.audubon.org/important-bird-areas

111. Wilcove DS, Rothstein D, Dubow J, Phillips A, Losos E. Quantifying threats to imperiled species in the United States. Bioscience. 1998;48: 607–615. doi:10.2307/1313420

112. Kansas Depoartment of Wildlife Parks and Tourism. Threatened and endangered wildlife. 2015 [cited 20 Feb 2015]. Available: http://kdwpt.state.ks.us/Services/Threatened-and-Endangered-Wildlife

113. U.S Fish and Wildlife Service. Least tern essential habitat areas in Kansas and Oklahoma. Recover plan Inter Popul least tern (Sterna antillarum). 1990. Available: http://ecos.fws.gov

114. Masters RE, Skeen JE, Garner JA. Red-cockaded woodpecker in Oklahoma: an update of Wood’s 1974-77 study. Proc Oklahoma Acad Sci. 1989;69: 27–31.

115. Laurencio LR, Fitzgerald LA. Atlas of distribution and habitat of the dunes sagebrush lizard (Sceloporus arenicolus) in New Mexico. Collge Station; 2010. Available: http://www.bison-m.org/documents/25024_LaurencioandFitzgerald2010.pdf

116. Colorado Parks and Wildlife. Preble’s meadow jumping mouse occupied range. Fort Collins; 2011.

117. Kansas Biological Survey. Kansas natural resource planner occurrence records for threatened and endangered species. 2015. Available: http://kars.ku.edu/maps/naturalresourceplanner/

118. Oklahoma Department of Wildlife Conservation. Oklahoma’s threatened and endangered species. 2015 [cited 15 Feb 2015]. Available: http://wildlifedepartment.com/wildlifemgmt/endangeredspecies.htm

119. Diamond D. Range-wide modeling of golden-cheeked warbler habitat (classes 2 and 3). Final Rep to Texas Park Wildl Dep. Austin; 2007. Available: https://tpwd.texas.gov/business/grants/wildlife/section-6/docs/birds/e72_final_report.pdf

120. Colorado Parks and Wildlife. Least tern foraging and production areas. Fort Collins; 2015.

121. U.S Fish and Wildlife Service. American burying beetle conservation priority areas shapefile and metadata. 2014. Available: http://fws.gov/southwest/es/oklahoma

122. Montana Natural Heritage Program. Occurrence data for terrestrial threatened and endangered species. Helena; 2018.

123. Texas Natural Diversity Database. American burying beetle, aplomado falcom, dunes sagebrush lizard, least tern, ocelot, and red-cockaded woodpecker element occurrence data exports. Austin; 2018.

124. Sawyer H, Nielson RM, Lindzey F, McDonald LL. Winter habitat selection of mule deer before and during development of a natural gas field. J Wildl Manage. 2006;70: 396–403.

125. Sawyer H, Kauffman MJ, Nielson RM. Influence of Well Pad Activity on Winter Habitat Selection Patterns of Mule Deer. J Wildl Manage. 2009;73: 1052–1061. doi:10.2193/2008-478

126. Wyoming Game and Fish Department. Recommendations for development of oil and gas resources within important wildlife habitats. Cheyenne; 2010. Available: https://www.nrc.gov/docs/ML1108/ML110810642.pdf

127. Vore J. Big game winter range recommendations for subdivision development in Montana: justification and rationale. Mont Fish, Wildl Park Prof Pap. 2012. Available: http://fwp.mt.gov

128. Taylor KL, Beck JL, Huzurbazar S V. Factors influencing winter mortality risk for pronghorn exposed to wind energy development. Rangel Ecol Manag. 2016;69: 108–116. doi:10.1016/j.rama.2015.12.003

129. Montana Fish Wildlife and Parks. Big game winter range habitat. 2010. Available: http://fwp.mt.gov/fishAndWildlife/conservationInAction/crucialAreas.html

130. Wyoming Game and Fish Department. Mule deer and pronghorn crucial winter ranges. Cheyenne; 2011.

131. Colorado Parks and Wildlife. Mule deer and pronghorn severe winter ranges. Fort Collins; 2015.

132. Western Electricity Coordinating Council. WECC environmental data viewer. 2018 [cited 2 Jul 2017]. Available: http://www.wecc.org

133. Ewert DN, Cole JB, Grmam E. Wind energy: Great Lakes regional guidelines. Rep to Nat Conserv Lansing, Michigan. 2011. Available: http://www.conservationgateway.org

134. Grodsky SM, Jennelle CS, Drake D. Bird mortality at a wind-energy facility near a wetland of international importance. Condor. 2013;115: 700–711. doi:10.1525/cond.2013.120167

135. Playa Lakes Joint Venture. Energy development siting recommendations for playas. 2017. Available: http://pljv.org/documents/PLJV_Energy_Development_Siting_Recommendations_Playas.pdf

136. Minnesota Department of Natural Resources. Wild, scenic, and recreational rivers. 2018. Available: https://gisdata.mn.gov

137. Western Hemisphere Shorebird Reserve Network. WHSRN sites. 2019 [cited 10 May 2019]. Available: http://www.whsrn.org

138. U.S Geological Survey. Protected areas database of the United States version 1.4. 2016. Available: https://www.usgs.gov/core-science-systems/science-analytics-and-synthesis/gap

139. Montana Natural Heritage Program. Wetlands of special significance. Helena; 2016.

140. Montana Natural Heritage Program. Species occurrence data for trumpter swan. Helena; 2018.

141. Argonne National Laboratory. West-wide wind mapping project: BLM-administered lands excluded from wind energy development. 2016. Available: http://www.mp.anl.gov

142. Comer PJ, Hak. J, Kindscher K, Muldavin E. Continent-scale landscape conservation design for temperate grasslands of the Great Plains and Chihuahuan Desert. Nat Areas J. 2018;38: 196–211.

143. Theobald DM. A general model to quantify ecological integrity for landscape assessments and U.S application. Landsc Ecol. 2013;28: 1859–1874. doi:10.1007/s10980-013-9941-6

144. Arkansas Oil & Gas Commission. Arkansas oil and gas wells. 2014 [cited 25 Oct 2018]. Available: https://gis.arkansas.gov

145. FracTracker Alliance. U.S oil and gas activity [for Illinois, Indiana, and Ohio]. 2016. Available: https://www.fractracker.org

146. Colorado Oil & Gas Conservation Commission. Well surface locations. 2018. Available: https://cogcc.state.co.us

147. Kansas Geological Survey. Oil and gas wells shapefile. 2018. Available: http://www.kgs.ku.edu/PRS/petroDB.html

148. Missouri Department of Natural Resources. Oil and gas wells tabular data. 2018. Available: https://dnr.mo.gov/geology/geosrv/oilandgas.htm

149. Montana Department of Natural Resources & Conservation. Oil and gas wells shapefile. 2018. Available: http://dnrc.mt.gov

150. Nebraska Oil and Gas Conservation Commission. Wells GIS shapefile. 2018. Available: http://www.nogcc.ne.gov/NOGCCPublications.aspx

151. New Mexico Energy Minerals and Natural Resources Department. Oil and gas shapefile. 2018. Available: http://www.emnrd.state.nm.us/OCD/ocdgis.html

152. North Dakota Department of Mineral Resources. Wells shapefile. 2018. Available: https://www.dmr.nd.gov/oilgas/

153. South Dakota Department of Environment and Natural Resources. Records of oil and gas drilling. 2018. Available: http://www.sdgs.usd.edu/SDOIL/oilgas_databases.aspx

154. Texas Railroad Commission. Oil and gas wells. Austin; 2018.

155. Wyoming Oil and Gas Conservation Commission. All wells tabular data. 2018. Available: http://pipeline.wyo.gov

156. Ostlie W. Untilled landscapes of the Great Plains. Minneapolis, Minnesota; 2003.

157. The Conservation Fund. Midwest wind energy HCP green infrastructure network. 2018. Available: http://tcfmidwestwind.conservationgis.com

158. The Nature Conservancy. Ecoregional conservation in the Osage Plains/Flint Hills Prairie. Minneapolis, Minnesota; 2000.

159. Colorado Natural Heritage Program. Level 3 potential conservation areas. Fort Collins; 2017.

160. Iowa Department of Natural Resources. Wind farm review layer. Des Moines; 2018.

161. Minnesota Department of Natural Resources. MBS sites of biodiversity significance. 2015. Available: https://gisdata.mn.gov

162. Minnesota Prairie Plan Working Group. Minnesota prairie conservation plan core areas, corridors, matirx habitat complexes, and strategic habitat complexes. St. Paul; 2017.

163. The Nature Conservancy. Gulf Coast Prairies and Marshes ecoregional conservation plan. San Antonio, Texas; 2002.

164. U.S Fish and Wildlife Service. Draft land protection plan: Austin’s Woods. 2012. Available: https://www.fws.gov/southwest/

165. U.S Department of Transportation. National transportation atlas database airport runways. 2017. Available: http://osav-usopendata.arcgis.com

166. Federal Aviation Administration. Airspace, special use airspace, and temporary flight restrictions. 2010 [cited 9 Dec 2015]. Available: https://www.faasafety.gov

167. Natural Resources Defense Council, U.S Department of Defense. Working with the Department of Defense: siting renewable energy development. 2013. Available: http://docs.nrdc.org/nuclear/files/nuc_13112001a.pdf

168. Federal Aviation Administration. U.S special use airspace shapefile. 2017. Available: http://ais-faa.opendata.arcgis.com

169. Federal Aviation Administration. Visual flight reference raster charts for Albuquerque, Billings, Brownsville, Cheyenne, Chicago, Cincinnati, Dallas-Ft. Worth, Denver, Detroit, El Paso, Great Falls, Green Bay, Houston, Kansas City, Memphis, Omaha, Salt Lake City, San Antonio, St. Louis,. 2017. Available: https://www.faa.gov/air_traffic/flight_info/aeronav

170. Vogt RJ, Ciardi EJ, Guenther RG. New criteria for evaluating wind turbine impacts on NEXRAD weather radars. Norman, Oklahoma; 2013. doi:004

171. National Oceanic and Atmospheric Administration. Climate data online radar mapping tool. 2017 [cited 31 Dec 2017]. Available: https://gis.ncdc.noaa.gov/map/viewer/#app=cdo

172. Federal Aviation Administration. DoD preliminary screening tool. 2018 [cited 3 Feb 2018]. Available: https://www.oeaaa.faa.gov

173. Federal Aviation Administration. Digital obstacle file. 2018. Available: https://www.faa.gov/air_traffic_flight_info/digital_products/dof

174. U.S Geological Survey. National elevation dataset. 2017. Available: http://ned.usgs.gov

175. AWS Truepower and National Renewable Energy Laboratory. Utility-scale land-based 80m wind maps for the United States. 2010 [cited 4 Dec 2013]. Available: http://apps2.eere.energy.gov/wind/windexchange/wind_maps.asp

176. Bailey BH, McDonald SL, Bernadett DW, Markus MJ, Elsohlz KV. Wind resource assessment handbook: fundamentals for conducting a successful monitoring program. Rep Prep Natl Renew Energy Lab. Albany, New York; 1997. Available: http://www.nrel.gov/docs/legosti/fy97/22223.pdf

177. Tennis MW, Clemmer S, Howland J. Assessing wind resources: a guide for landowners, project developers, and power suppliers. Union Concerned Sci Brief Pap. Cambridge, Massachusetts; 1999. Available: http://www.ucsusa.org/sites/default/files/legacy/assts/documents/clean_energy/wind_resource_assessment.pdf

178. Langreder W. Wind resource and site assessment. In: Tong W, editor. Wind power generation and wind turbine design. Billerica, Massachusetts: WIT Press; 2010. pp. 49–87.

179. Hau E, von Renouard H. The wind resource. Wind turbines: fundamentals, technologies, application, economics. Berlin: Springer Science & Business Media; 2013. pp. 505–548.

180. White T, Kuba J, Thomas J. Data driven generation siting for renewables integration in transmission planning. Proc ESRI User Conf July 14-18, 2014, San Diego, Calif. 2014.

181. Oklahoma Legislature. Oklahoma Statutes, Title 17, § 160.20. 2017. doi:10.1145/3132847.3132886

182. Oklahoma State Department of Education. Oklahoma public schools 2014-2015 shapefile. 2015. Available: http://okmaps.org

183. U.S Geological Survey. 2017 geographic names information system text file for Oklahoma. 2017. Available: http://viewer.nationalmap.gov

184. Rothschild S. Flint Hills may nbot get wind of power plan. Lawrence Journal World. 15 Jan 2005: 1b, 3b. Available: https://www2.ljworld.com/news/2005/jan/15/flint_hills_may/

185. Kansas Biological Survey. Kansas natural resource planner Tallgrass Heartland shapefile. 2015. Available: http://kars.ku.edu/maps/naturalresourceplanner/

186. Illinois General Assembly. Illinois Statutes 525 ILCS 30/1-26. 2018.

187. Prairie State Conservation Coalition. Illinois protected natural lands (Illinois Nature Preserves Commission, Illinois Natural Areas Inventory, and Illinois Department of Natural Resources properties). Springfield; 2015.

188. Minnesota Department of Natural Resources. Minnesota trails: division of parks & trails. 2016. Available: https://gisdata.mn.gov

189. Minnesota Department of Natural Resources. Guidance for commerical wind energy projects. 2018. Available: https://www.dnr.state.mn.us

190. Denholm P, Jackson M, Ong S, Hand M. Land-use requirements of modern wind power plants in the United States. Technical report NREL/TP-6A2-45834. Golden, Colorado; 2009.

191. American Wind Energy Association. U.S wind industry third quarter 2019 market report. Washington, D.C.; 2019.

192. U.S Department of Energy. 2017 wind technologies market report. 2017. Available: http://eta-publications.lbl.gov

193. Oteri F, Baranowski R, Baring-gould I, Tegen S. 2017 state of wind development in the United States by region. Technical report NREL/TP-5000-70738. Golden, Colorado; 2018.

194. Janke AK, Anteau MJ, Stafford JD. Prairie wetlands confer consistent migrant refueling conditions across a gradient of agricultural land use intensities. Biol Conserv. 2019;229: 99–112. doi:10.1016/j.biocon.2018.11.021

195. Hayes MA, Cryan PM, Wunder MB. Seasonally-dynamic presence-only species distribution models for a cryptic migratory bat impacted by wind energy development. PLoS One. 2015;10: 1–20. doi:10.1371/journal.pone.0132599

196. Arnett EB, Johnson GD, Erickson WP, Hein CD. A synthesis of operational mitigation studies to reduce bat fatalities at wind energy facilities in North America. A Rep Submitt to Natl Renew Energy Lab. 2013. Available: http://www.batsandwind.org

197. Arnett EB, Baerwald EF, Mathews F, Rodrigues L, Rodríguez-Durán A, Rydell J, et al. Impacts of wind energy development on bats: a global perspective. In: Voigt CC, Kingston T, editors. Bats in the anthropocene: conservation of bats in a changing world. Cham, Switzerland: Springer International Publishing; 2015. pp. 295–323.

198. Hayes MA, Hooton LA, Gilland KL, Grandgent C, Smith RL, Lindsay SR, et al. A smart curtailment approach for reducing bat fatalities and curtailment time at wind energy facilities. Ecol Appl. 2019;29: 1–18. doi:10.1002/eap.1881

199. Cryan PM, Gorresen PM, Hein CD, Schirmacher MR, Diehl RH, Huso MM, et al. Behavior of bats at wind turbines. Proc Natl Acad Sci U S A. 2014;111: 15126–15131. doi:10.1073/pnas.1406672111

200. Arnett EB, Huso MMP, Schirmacher MR, Hayes JP. Altering turbine speed reduces bat mortality at wind-energy facilities. Front Ecol Environ. 2011;9: 209–214. doi:10.1890/100103

201. Martin CM, Arnett EB, Stevens RD, Wallace MC. Reducing bat fatalities at wind facilities while improving the economic efficiency of operational mitigation. J Mammal. 2017;98: 378–385. doi:10.1093/jmammal/gyx005

202. Arnett EB, Hein CD, Schirmacher MR, Huso MMP, Szewczak JM. Evaluating the effectiveness of an ultrasonic acoustic deterrent for reducing bat fatalities at wind turbines. PLoS One. 2013;8: 1–11. doi:10.1371/journal.pone.0065794

203. Weller TJ, Castle KT, Liechti F, Hein CD, Schirmacher MR, Cryan PM. First direct evidence of long-distance seasonal movements and hibernation in a migratory bat. Sci Rep. 2016;6: 1–7. doi:10.1038/srep34585

